# HPV status and oxygen tension shape transcriptomic, inflammatory, and cell cycle responses in HNSCC treated with ionizing radiation

**DOI:** 10.1101/2025.10.26.684626

**Authors:** Jana Pereckova, Filip Zavadil Kokas, Simona Voznicova, Ondrej Vasicek, Jitka Holcakova, Roman Hrstka, Tomas Perecko

## Abstract

Head and neck squamous cell carcinoma (HNSCC) comprises biologically distinct subtypes defined by human papillomavirus (HPV) status, which may influence treatment outcomes. Hypoxia is a common feature of solid tumors, including HNSCC, and can reduce radiotherapy efficacy by modulating DNA repair, cell-cycle progression, and inflammatory signaling. However, the combined impact of hypoxia and HPV status on cellular radiosensitivity remains poorly defined. We investigated the influence of hypoxia (1% O_2_) and gamma-irradiation on proliferation, cell cycle, apoptosis, transcriptomic profiles, and cytokine/chemokine secretion in three HNSCC cell lines: HPV-negative FaDu and Detroit-562 (metastatic origin), and HPV-positive 2A3. Cells were cultured under chronic hypoxic or ambient conditions, with or without irradiation. Independent of HPV status or metastatic phenotype, hypoxia prolonged the cell doubling time, while only irradiated cells (under both oxygen conditions) exhibited features of an intermediate epithelial–mesenchymal transition phenotype. Treatment of HPV-positive cells significantly decreased the number of upregulated genes, reduced cytokine secretion (IL-8, SERPINE1), and affected all phases of the cell cycle compared with HPV-negative cells. Furthermore, only the metastatic cell line showed no change in cleaved caspase-3 levels after irradiation in hypoxia and produced higher levels of cytokines (IL-8, SERPINE1). Our findings highlight how tumor oxygenation and HPV status intersect to modulate the radiation response in HNSCC. These insights may guide the development of personalized, biomarker-driven radiotherapy strategies that account for both the tumor microenvironment and viral etiology.

## Introduction

Hypoxia within the tumor microenvironment promotes tumor development, progression, and metastasis, and is associated with multiple factors that influence the response to anticancer treatments, including radiotherapy (reviewed in (1)). Tumor cell radiosensitivity is known to decrease significantly under hypoxic conditions (2). Long-term exposure of cells to moderate hypoxia prior to irradiation has been shown to result in pronounced, cell line-specific differences in radiation response, indicating distinct mechanisms of hypoxia adaptation (3).

Head and neck squamous cell carcinoma (HNSCC) ranks among the ten most common cancers worldwide (4), and approximately 80% of patients with HNSCC have an indication for radiotherapy (5). HNSCC tumors are commonly characterized by hypoxia, which contributes to poor prognosis and reduced responsiveness to radiotherapy (reviewed in (6)). HNSCC can be classified by human papillomavirus (HPV) status: HPV-negative tumors are generally associated with tobacco or alcohol abuse, whereas HPV-positive tumors are primarily linked to HPV-16 infection (6). The carcinogenic effects of HPV are primarily mediated by the E6 and E7 oncoproteins, which inhibit the functions of p53 and the retinoblastoma tumor suppressor protein, respectively (7). HPV status may influence several biological characteristics of HNSCC. Notably, HPV-positive HNSCC cells exhibit a pronounced G2/M arrest following irradiation, reflecting impaired DNA double-strand break repair rather than enhanced apoptosis or a functional p53 response (8). Although HPV-positive cells generally demonstrate increased radiosensitivity compared to HPV-negative cells, they exhibit similar relative radioresistance under hypoxic conditions (9). This underscores the need for integrated analyses to guide personalized radiotherapy strategies.

Other components of the HNSCC tumor microenvironment, such as cytokines and chemokines, may reflect disease status, treatment response, or intrinsic (e.g., HPV status) and extrinsic (e.g., hypoxia) characteristics of the tumor cells (10). Several cytokines have been reported to be elevated in patients with HNSCC—such as interleukin-8 (IL-8), vascular endothelial growth factor, and macrophage migration inhibitory factor (MIF), whereas others have been described as decreased (e.g., IL-2, IL-4, Serpin E1, GROα) or unchanged (e.g., G-CSF, IFN-γ, TNFα) when compared with healthy controls. Notably, findings for certain cytokines, including IL-6, IL-8, and Serpin E1, have been inconsistent across studies (10-13).

Despite these well-established findings, many mechanistic studies on cancer cells *in vitro* are still conducted under ambient air conditions. The objective of our study was to compare the responses of HNSCC cell lines to gamma-irradiation under ambient and hypoxic conditions, as well as between primary and metastatic origins, taking HPV status into account. We focused on assessing proliferation, cell cycle progression, apoptosis, and cytokine/chemokine production in response to radiation and oxygen. Additionally, we performed transcriptomic profiling to identify condition-specific gene expression signatures associated with hypoxia and radiation response in two isogenic HNSCC cell lines. These findings may help personalize radiotherapy according to hypoxia status.

## Results

### Transcriptomic responses to hypoxia and irradiation in HNSCC

The total numbers of differentially expressed genes (DEGs) in isogenic HNSCC cell lines FaDu and 2A3 detected under each condition are summarized in Figure 1A–D, while volcano plots (Figure S1) illustrate the magnitude and direction of transcriptional changes. DEGs in FaDu and 2A3 cells under hypoxic conditions revealed distinct adaptive responses (Figure 1E). This reflects a coordinated transcriptional program aimed at enhancing survival, promoting tissue remodeling, and enabling communication within a stressed microenvironment. In contrast, downregulated genes in FaDu and 2A3 cell lines were predominantly involved in biosynthetic, metabolic, and structural processes (Figure 1E). Upon combined exposure to hypoxia and gamma-irradiation, FaDu and 2A3 cells exhibited cell line-specific transcriptional responses (Figure 1E). In FaDu cells, upregulated genes were enriched in pathways related to synaptic signaling, neurogenesis, and cell adhesion, potentially reflecting stress-induced plasticity and neural-like reprogramming, a phenomenon observed in epithelial tumors following genotoxic stress and often associated with treatment resistance. In contrast, 2A3 cells responded with a broad activation of morphogenetic and structural processes, including cytoplasmic translation, morphogenesis, migration, and neurogenesis suggesting a reparative or pro-survival shift possibly linked to radio-adaptive plasticity. Downregulated genes in FaDu included those involved in glucuronidation, pigment metabolism, and gland development, pointing to further suppression of differentiation and metabolic stability under compounded stress. Downregulated genes in 2A3 were associated with daunorubicin metabolism, collagen catabolism, intermediate filament organization, and axon development, indicating suppression of matrix remodeling and certain structural pathways.

**Figure 1.**
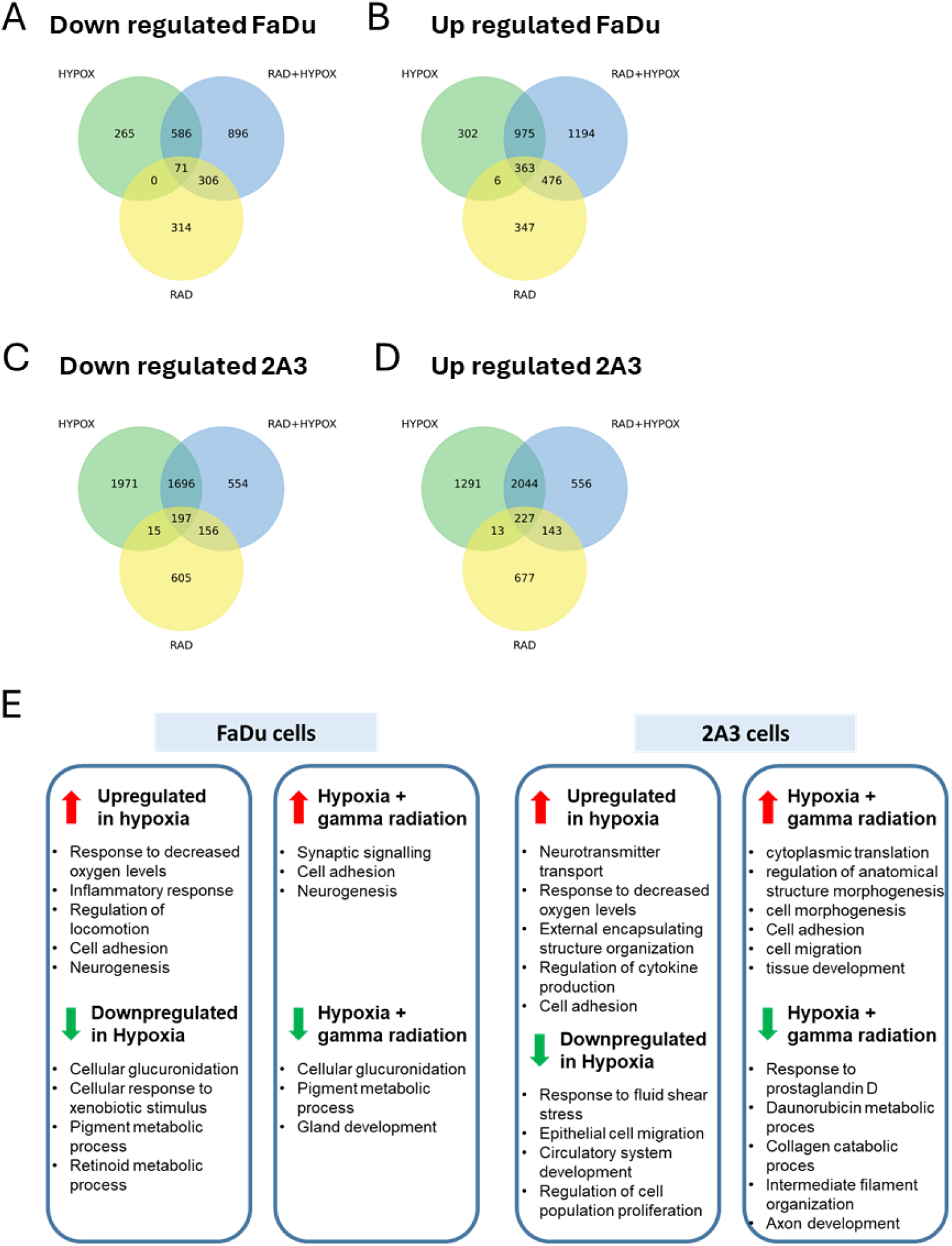
Transcriptomic responses to hypoxia and irradiation in HNSCC. (A-D) Venn diagrams of differentially expressed genes in FaDu (A-B) and 2A3 (C-D) cell lines. Venn diagrams illustrate the number of significantly regulated genes under three conditions: hypoxia (green), irradiation (yellow), and the combination of hypoxia and irradiation (blue). Numbers in the overlapping areas indicate genes commonly regulated under the corresponding conditions. Genes were considered differentially expressed if they had an adjusted p-value (padj) <0.1. (E) Biological processes differentially regulated in FaDu and 2A3 HNSCC cell lines under hypoxia and combined hypoxia with gamma irradiation. The figure illustrates biological processes significantly upregulated (red) or downregulated (green) in FaDu and 2A3 cells in response to hypoxia alone or in combination with gamma radiation. The analysis was performed using the WebGestalt 2024 platform and included only genes that were consistently detected across all three independent biological replicates and met the inclusion criteria of adjusted p < 0.1 and |log_2_FC| ≥ 2.

### Effects of hypoxia on HNSCC cell proliferation

Under normoxia, 2A3 cells (Figure 2A) showed a baseline doubling times (DT) comparable to Detroit-562. The shorter DT of FaDu cells suggest higher proliferative activity. Next, we assessed the growth kinetics of HNSCC cell lines under hypoxia. As shown in Figure 2A, all lines showed increased DT under hypoxia, with FaDu and Detroit-562 exhibiting significantly delayed proliferation. In contrast, 2A3 cells showed no significant change in DT but displayed growth exhaustion, failing to proliferate beyond two weeks during prolonged hypoxia, whereas FaDu and Detroit-562 maintained stable growth. Consequently, 2A3 cells were cultured under hypoxia for up to 14 days and re-established from ambient conditions before each new experiment.

**Figure 2.**
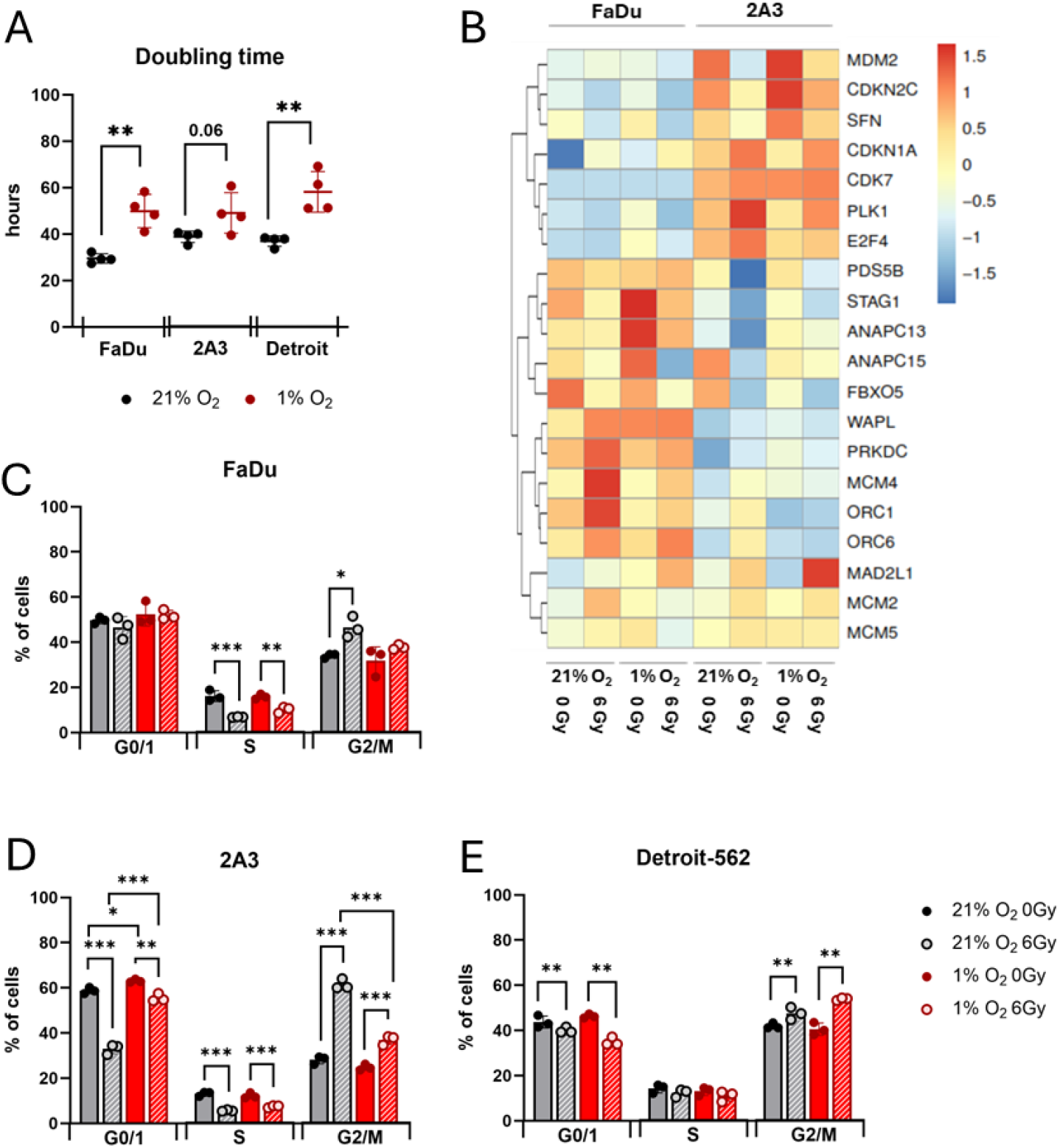
Proliferation and cell cycle regulation under different oxygen conditions and following gamma-irradiation. (A) FaDu, 2A3, and Detroit-562 cell lines were cultured under normoxic (21% O_2_) or hypoxic (1% O_2_) conditions. Cell numbers were determined 48 and 72 hours after plating, and doubling times were calculated as described in Methods. Data represent the mean ± SD from four independent experiments (n = 4). Statistical analysis was performed using an unpaired t-test (**p < 0.01). (B) Heatmap of the top 20 DEGs. Averaged data from three independent experiments are shown. The color bar is showing the values of z-score for each gene after library size normalization via DESeq2 software. Cell cycle distribution was analyzed 24 hours after 6 Gy gamma-irradiation in (C) FaDu, (D) 2A3, and (E) Detroit-562 cells cultured under normoxic or hypoxic (1% O_2_) conditions. Results are presented as the percentage of cells in G0/1, S, or G2/M phases. Data represent the mean ± SD from three independent experiments (n = 3). Statistical analysis was performed using two-way ANOVA with Tukey’s multiple comparisons test; *p < 0.05, **p < 0.01, ***p < 0.001.

### Combined effects of hypoxia and gamma-irradiation on cell cycle progression in HNSCC

Next, we analyzed cell cycle distribution 24 hours after gamma-irradiation to assess simultaneous effects of hypoxia and radiation (Figure 2B-E). Based on the transcriptomic analysis, 2A3 cells— unlike FaDu—showed, under hypoxia, increased expression of cell-cycle negative regulators (*MDM2, CDKN2C*) and decreased expression of the positive regulator *ANAPC15*. In contrast, FaDu showed increased expression of *STAG1, ANAPC13/15*, and *WAPL*, which are positive regulators of the cell cycle. Gamma-irradiation under ambient conditions differentially affected the DEGs in both cell lines: in FaDu, we observed increased expression of *WAPL, PRKDC, MCM4*, and *ORC1/6*, and the effects of irradiation on these genes were mitigated under hypoxia, except for ORC6; in 2A3 up-regulation of *CDK7, PLK1*, and *E2F4* was detected. On the other hand, *PDS5B, STAG1*, and *ANAPC13/15* were downregulated in 2A3 under the same conditions. This effect was less evident in 2A3 irradiated under hypoxia.

Based on flowcytometry analysis, FaDu cells showed no change in the G0/G1 phase under either condition (Figure 2C), whereas Detroit-562 cells exhibited a decreased G0/G1 fraction after irradiation, independent of oxygen (Figure 2E). In contrast, 2A3 cells were the most responsive: irradiation reduced G0/G1 accumulation, hypoxia increased it, and hypoxia significantly mitigated this irradiation-induced decrease (Figure 2D). Irradiated FaDu and 2A3 cells displayed oxygen-independent S-phase depletion, unlike Detroit-562. All cell lines showed significant G2/M arrest after irradiation under normoxia (Figure 2C–E), most pronounced in 2A3. Under hypoxia, Detroit-562 and 2A3 also exhibited G2/M arrest, but only 2A3 showed a significant hypoxia-induced reduction.

### Impact of hypoxia and gamma-irradiation on apoptosis in HNSCC

We next examined whether hypoxia or gamma-irradiation affected apoptosis. Forty-eight hours after irradiation, cleaved caspase-3 levels increased under ambient conditions (Figure 3A). In FaDu and 2A3 cells, this increase was significant under both normoxia and hypoxia, whereas in Detroit-562, cleaved caspase-3 was not elevated under hypoxia compared to normoxia (Figure 3A).

**Figure 3.**
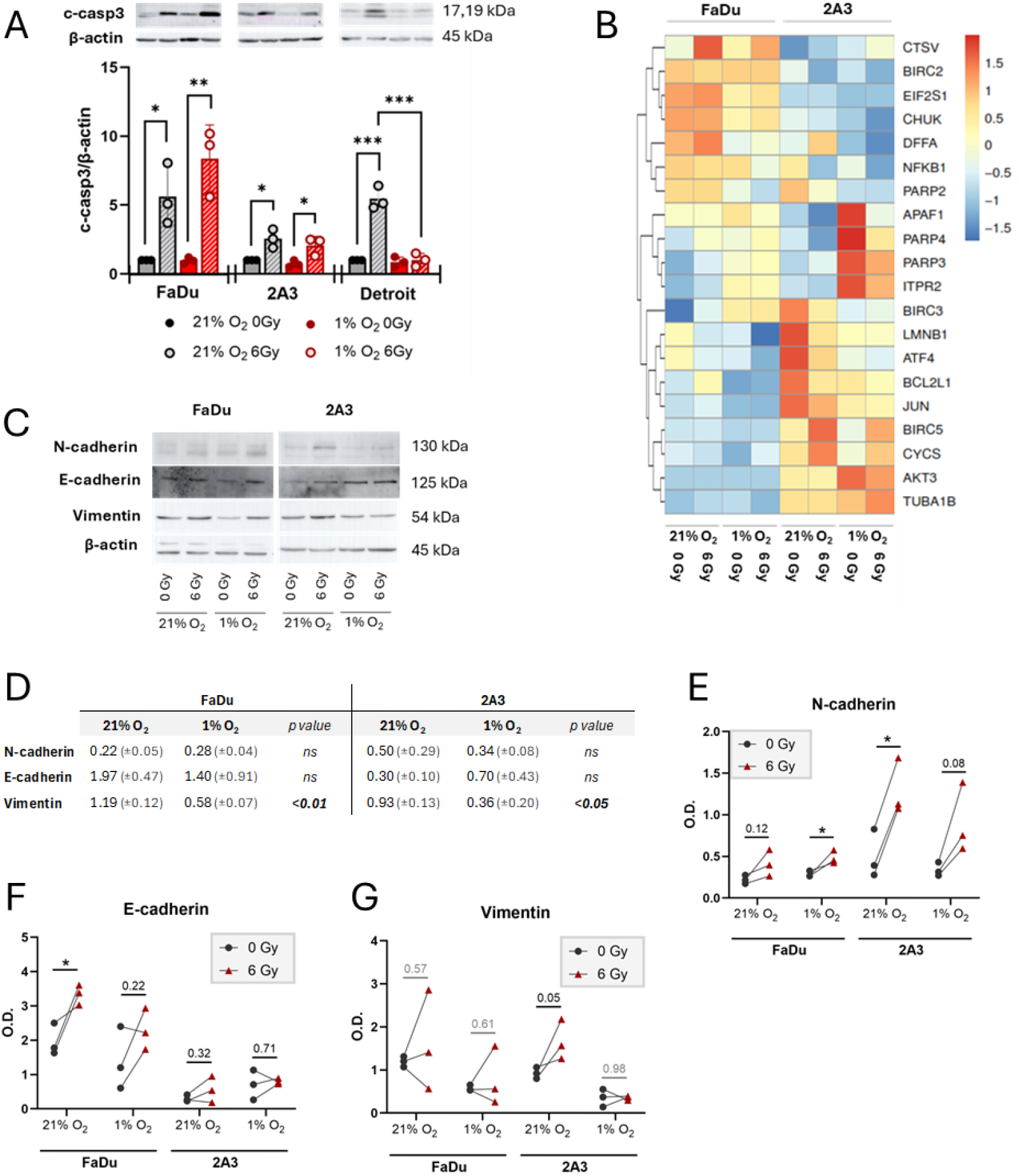
Effects of hypoxia and gamma-irradiation on apoptosis and EMT in HNSCC. (A) Cleaved caspase-3 levels in FaDu, 2A3, and Detroit-562 cells cultured under normoxic (21% O_2_) or hypoxic (1% O_2_) conditions, assessed 48 hours after 6 Gy gamma-irradiation. Results are shown as fold change relative to non-irradiated normoxic controls. Data represent mean ± SD from three independent experiments (n = 3). Statistical analysis: two-way ANOVA with Tukey’s multiple comparisons test; *p < 0.05, **p < 0.01, ***p < 0.001. (B) Heatmap of the top 20 DEGs. Averaged data from three independent experiments are shown. The color bar is showing the values of z-score for each gene after library size normalization via DESeq2 software. (C) Representative western blots of N-cadherin, E-cadherin, and vimentin in FaDu and 2A3 cells. (D) Quantification of protein levels expressed as mean optical density (O.D.) ± SD, normalized to β-actin, in FaDu and 2A3 cells under normoxic or hypoxic conditions. (E–G) Effects of hypoxia and irradiation on N-cadherin, E-cadherin, and vimentin levels in FaDu and 2A3 cells under normoxic and hypoxic conditions. Data represent mean ± SD from three independent experiments (n = 3). Statistical analysis: unpaired t-test (*p < 0.05).

Transcriptomic analysis (Figure 3B) revealed that FaDu cells have lower baseline expression of anti-apoptotic genes (*BIRC3/5, BCL2L1* and *AKT3*) and higher expression of pro-apoptotic effectors (*EIF2S1, DFFA, PARP2* and *APAF-1*) compared to 2A3. Under hypoxia (1% O_2_), FaDu exhibited downregulation of pro-apoptotic mediators (*DFFA, CYCS, PARP2*), while most anti-apoptotic genes remained unchanged (*BIRC2/5, AKT3*). On the other hand, 2A3 showed decreased expression of both anti/pro-apoptotic genes (*BIRC3, BCL2L1*, and *DFFA, CYCS*). Following irradiation under normoxia, FaDu showed upregulation of pro-apoptotic genes (*EIF2S1, DFFA*). In 2A3 cells, irradiation downregulated anti-apoptotic genes (*BIRC2/3, BCL2L1*). These effects were attenuated under hypoxia.

### Hypoxia and gamma-irradiation and EMT in HNSCC

To observe how hypoxia and irradiation could change the metastatic potential of the cells, we analyzed genes and proteins associated with epithelial mesenchymal transition (EMT). Hypoxia significantly reduced vimentin in FaDu and 2A3, while N-cadherin and E-cadherin remained unchanged (Figure 3C, Figure D). Irradiation further modulated EMT markers (Figure 3E–G): N-cadherin increased in FaDu under hypoxia and in 2A3 under normoxia; E-cadherin was upregulated in FaDu under normoxia; and vimentin showed variable responses, with 2A3 trending toward upregulation. Overall, hypoxia primarily downregulated vimentin level, while irradiation induced N-cadherin and E-cadherin expression in FaDu, indicating cell line–specific EMT modulation with potential implications for invasiveness.

### Cytokine profiling and ELISA quantification

To identify cytokines and chemokines modulated by hypoxia or gamma-irradiation in HNSCC cell lines, we analyzed our transcriptomic data (Figure 4A). In FaDu cells, hypoxia downregulated *CSF1, CSF1R*, and *CSF2RA*, indicating reduced GM-CSF signaling, and upregulated *IL24* and *CXCR6*, favoring immune activation and potential tumor suppression. Upon irradiation under hypoxia, FaDu maintained elevated *IL24* and further increased *CXCR6*, with only modest *CSF2RA* induction and a small rise in invasion markers such as *CCL26* and *GDF11*, suggesting a shift toward a more immune-permissive yet more aggressive phenotype. In 2A3 cells, hypoxia alone induced *CXCR6* and also upregulated *CSF1, EPOR*, and *CCL26*, supporting a more pro-tumorigenic, invasive profile. Under hypoxic irradiation, 2A3 partially reversed this progression-associated profile, with reductions in *CSF1, EPOR*, and *CCL26*, indicating partial normalization of the hypoxia-driven invasive signature.

**Figure 4.**
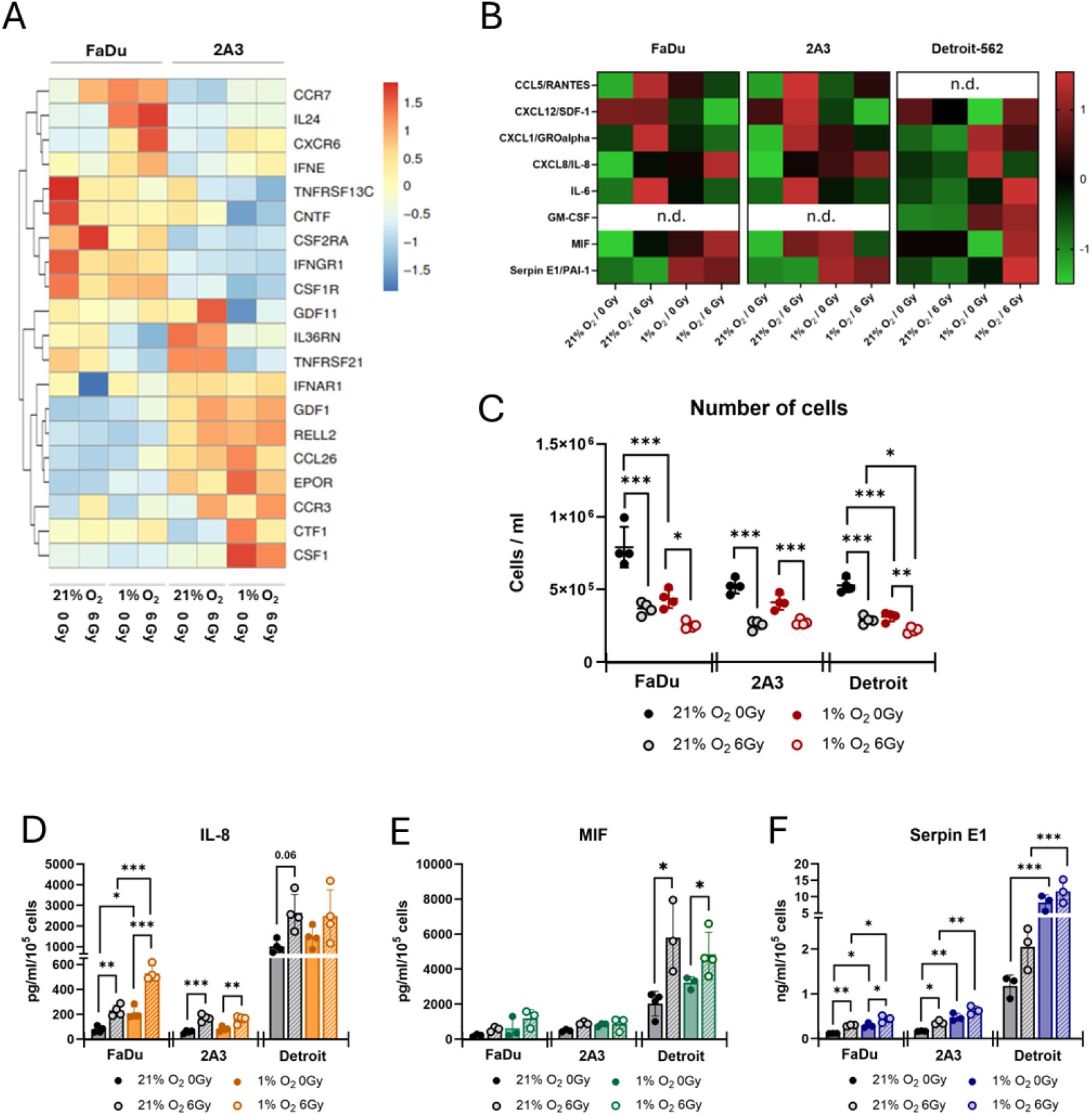
Effects of hypoxia and gamma-irradiation on cytokine/chemokine release in HNSCC. (A) Heatmap of the top 20 DEGs. Averaged data from three independent experiments are shown. The color bar is showing the values of z-score for each gene after library size normalization via DESeq2 software. (B) Cytokine/chemokine production in response to hypoxia, gamma-irradiation, and their combination. Conditioned media from FaDu, 2A3, and Detroit-562 cells were collected 48 hours after gamma-irradiation under normoxic or hypoxic conditions. Media from three independent experiments per cell line were pooled for analysis. Results are presented as z-scores of log_10_-transformed normalized data. n.d. = not detected. (C) Cell numbers of FaDu, 2A3, and Detroit-562 cultured under normoxic (21% O_2_) or hypoxic (1% O_2_) conditions, assessed 48 hours after 6 Gy gamma-irradiation. Data represent mean ± SD from four independent experiments (n = 4). Statistical analysis: two-way ANOVA with Tukey’s multiple comparisons test; *p < 0.05, **p < 0.01, ***p < 0.001. Levels of IL-8 (D), MIF (E), and Serpin E1 (F) in supernatants from FaDu, 2A3, and Detroit-562 cells cultured under normoxic or hypoxic conditions, assessed 48 hours after gamma-irradiation. Concentrations were normalized to cell numbers per condition. Data represent mean ± SD from three to four independent experiments (n = 3–4). Statistical analysis: two-way ANOVA with Tukey’s multiple comparisons test; *p < 0.05, **p < 0.01, ***p < 0.001.

In both FaDu and 2A3 cells, irradiation downregulated canonical TGF-β components such as *TGFB2, TGFBR2, SP1*, and *SMAD3*, while simultaneously upregulating bone morphogenic protein (BMP) ligands (*BMP6, BMP8A, GDF11*) and TGF-β inhibitors (*SKI, ID3*). This pattern suggests a shift away from canonical TGF-β/SMAD signaling, which is often associated with EMT and tumor progression, toward BMP-driven or inhibitory networks potentially restricting EMT and favoring differentiation. Hypoxia alone further reinforced this trend, suppressing *TGFB1/2, SMAD1/5/6*, and *PITX2* while increasing *SMAD3, INHBA, FST*, and *SMURF2*, consistent with remodeling of TGF-β signaling and induction of antagonistic feedback loops. Strikingly, the combination of hypoxia and irradiation produced the most pronounced effects, synergistically upregulating *IL24, IFNE* in FaDu, and chemokine receptors (*CCR7, CXCR6* in FaDu, and *CCR3* in *2A3*), while strongly repressing *SP1, TGFB2, SMAD1/6*, and *PITX2* (Figures 4A, S2, S3).

As shown in cytokine array (Figure 4B), the cell lines differed in both detectable cytokine/chemokine profiles and responsiveness to hypoxia, irradiation, or their combination. FaDu and 2A3 produced CCL5/RANTES, which was absent in Detroit-562. In contrast, Detroit-562 secreted high levels of GM-CSF, not detected in the other two lines. Under normoxia, FaDu and 2A3 exhibited similar cytokine and chemokine induction after irradiation (Figure 4B), including increased CCL5/RANTES, CXCL12/SDF-1, CXCL1/GROα, CXCL8/IL-8, and IL-6. Under hypoxia, MIF and Serpin E1/PAI-1 were elevated, while Detroit-562 showed broader upregulation across nearly all detectable analytes. Based on these findings and signal-to-background ratios, IL-8, MIF, and Serpin E1/PAI-1 (hereafter Serpin E1) were selected for ELISA quantification.

For the ELISA experiments, we first assessed the effects of hypoxia and gamma-irradiation on cell numbers. A marked reduction was observed in non-irradiated hypoxic controls at 48 hours post-sham irradiation, with significant decreases in FaDu and Detroit-562 cells (Figure 4C). Similarly, 6 Gy irradiation significantly reduced cell counts in all cell lines, more strongly under normoxia, while irradiation under hypoxia caused a smaller yet significant decline. Among the three cell lines, only Detroit-562 showed a statistically significant difference between normoxic and hypoxic irradiation (Figure 4C).

IL-8 production in FaDu cells was dependent on both oxygen and radiation, with a significant interaction effect (Figure 4D). In contrast, 2A3 cell line exhibited a radiation-dependent increase in IL-8, with no significant effect of oxygen, a trend that was also observed in Detroit-562 cells (Figure 4D). Notably, IL-8 production in 2A3 was 2–3 times lower than in FaDu, whereas Detroit-562 cells produced IL-8 at levels approximately an order of magnitude higher than FaDu under comparable conditions (Figure 4D).

MIF levels were modestly elevated in FaDu and 2A3 cells in response to hypoxia or irradiation, although these changes did not reach statistical significance (Figure 4E). In contrast, gamma-irradiation significantly increased MIF production in Detroit-562 cells (Figure 4E). While FaDu and 2A3 cells exhibited comparable maximum MIF levels, Detroit-562 cells secreted 4-to 7-fold higher amounts under the same conditions (Figure 4E).

Serpin E1 levels increased significantly in response to hypoxia across all three HNSCC cell lines (Figure 4F). In FaDu and 2A3 cells, irradiation under ambient air conditions also led to a significant increase in Serpin E1 levels, an effect that was not observed in Detroit-562 cells. Notably, in both FaDu and 2A3, irradiation under hypoxic conditions resulted in a significantly greater additive effect compared to irradiation under normoxia.

IL-8, MIF, and Serpin E1 levels 24 hours after gamma-irradiation, illustrating time-dependent release patterns, are shown in Figure S4.

## Discussion

HPV-associated HNSCC constitute a biologically distinct subgroup with a generally favorable prognosis, yet the determinants of their differential treatment response—particularly under hypoxia—remain incompletely defined. Because hypoxia is prevalent in HNSCC and modulates radiobiology and inflammatory signaling, we examined how oxygen tension (ambient vs. 1% O_2_) shaped HPV-positive and HPV-negative responses to gamma-irradiation. To this end, we integrated analyses of proliferation, cell-cycle distribution, and apoptosis with transcriptomic profiling and cytokine/chemokine secretion to delineate mechanisms of treatment responsiveness.

Increased doubling time under hypoxia aligned with reports that hypoxia prolongs the lag phase of HNSCC proliferation, even if subsequent growth rates remain similar (14). Hypoxia slowed proliferation in all cell lines, but only 2A3 showed significant G0/G1 accumulation, indicating a distinct adaptation. In HeLa cells (HPV18+), chronic hypoxia reduced proliferation with G-phase arrest, suggesting that hypoxia blocks G–S transition (15), whereas in other HNSCC models, hypoxia induced cyclin D1 expression to promote G1–S transition (16). These alterations in cell-cycle progression are consistent with the transcriptional changes observed in our analyses, reflecting a broader metabolic and regulatory reprogramming aimed at conserving energy and supporting cell survival under oxygen limitation. In FaDu cells, suppression of energy-demanding pathways including glucuronidation, xenobiotic metabolism, and retinoid and pigment processing suggests a metabolic reprogramming in response to oxygen deprivation, in line with hypoxia-driven shifts mediated by HIF-1α (17). In 2A3 cells, downregulation of genes involved in epithelial dynamics and tissue development may reflect broader inhibition of morphogenetic programs under hypoxic stress (18). Overall, these findings illustrate how hypoxia drives the repression of energetically costly and differentiation-related pathways in a cell line-specific manner.

At baseline, HPV-negative and HPV-positive HNSCC cell lines, including 2A3, showed no differences in cell-cycle distribution (19, 20). Consistent with our results, gamma-irradiation induced G2/M arrest in both, with stronger and longer arrest in HPV-positive cells (19-22), though some reported this only in HPV-positive HNSCC (8). We also observed S-phase reduction in FaDu and 2A3 cells post-irradiation, irrespective of HPV status, contrasting Arenz et al., who reported it only in HPV-negative cells (21).

Few studies have examined radiation–cell cycle interactions under hypoxia in HNSCC. In other cancers, hypoxia can impair radiation-induced G2/M arrest (23). Ha et al. showed that gamma-irradiation failed to induce G2/M arrest under chemical hypoxia (24). Similarly, U87MG-E7 glioma cells lost G2/M arrest under hypoxia due to HPV16 E7 (25). Consistent with this, HPV-positive 2A3 cells showed reduced radiation-induced G2/M arrest under hypoxia with persistent G0/G1 accumulation. Detroit-562 cells, however, maintained G2/M arrest regardless of oxygen, suggesting resistance to hypoxia-induced modulation.

These findings underscore cell line–specific interactions between hypoxia, irradiation, and HPV status in shaping cell cycle responses. Such functional differences likely originate from underlying transcriptional adaptations, as our data reveal distinct molecular programs activated under combined hypoxia and irradiation. Specifically, gamma-irradiation under hypoxic conditions not only enhances the repression of energy-demanding pathways but also drives divergent transcriptional responses such as stress-induced plasticity in FaDu cells and morphogenetic activation in 2A3 cells, both of which may influence tumor progression and radiotherapy outcomes. Similar processes involving metabolic reprogramming and morphogenetic activation have also been reported in other epithelial malignancies e.g. pancreatic cancer, where they contribute to enhanced adaptability and invasive behavior (26).

In addition to cell cycle regulation, gamma-irradiation induced cleavage of caspase-3—a key effector caspase in both intrinsic and extrinsic apoptotic pathways—in all three HNSCC cell lines under ambient conditions. Notably, cleaved caspase-3 did not increase following irradiation under hypoxia in Detroit-562 cells, in contrast to FaDu and 2A3, which may reflect a shift toward checkpoint arrest rather than apoptosis (G2/M arrest preserved in hypoxia). Compared with HPV-negative cells, HPV-positive cells at ambient conditions showed increased caspase-3/7 activity after irradiation (19, 27). However, irradiation induced comparable apoptosis in FaDu and 2A3 (20). Long-term hypoxia was not associated with increased apoptosis in HPV-18–positive cancer cells after irradiation compared to normoxia (15). Interestingly, *in vivo*, the hypoxic fraction of HPV-positive tumors decreased after irradiation, an effect not observed in HPV-negative tumors (28). Therefore, the differential response of HPV-positive and HPV-negative HNSCC to irradiation under hypoxia highlights the potential of pharmacological interventions, such as AKT inhibition, to facilitate stratified treatment strategies (29). These findings also suggest that in some HNSCC subtypes, such as Detroit-562, hypoxia may divert irradiated cells from caspase-3–dependent apoptosis, warranting investigation of alternative pathways including senescence, lysosome-mediated cell death, autophagy, or mitotic catastrophe.

Although high pretreatment IL-8 has been associated with worse overall survival, it did not correlate with hypoxia staining (30). In our work, IL-8 increased under hypoxia only in FaDu cells, whereas irradiation upregulated IL-8 across all lines; a significant hypoxia–irradiation interaction was observed only in FaDu. Detroit-562 cells secreted markedly higher IL-8 than FaDu and 2A3. Prior studies reported both increases and decreases in IL-8 after irradiation, regardless of HPV status (31, 32). Clinically, a subgroup of HPV-positive patients with elevated pretreatment IL-8 showed improved outcomes compared with HPV-negative patients (10). However, baseline IL-8 was higher in patients who later progressed, irrespective of HPV status, and all progressing HPV-negative patients displayed elevations in this high-risk factor (33). These findings emphasize the need for combined hypoxia- and HPV-based stratification and suggest that integrating cytokine/chemokine risk profiles with HPV status may improve identification of patients at elevated risk of progression. Moreover, it appears that cytokine regulation under combined hypoxic and irradiative stress is not limited to IL-8, but reflects a wider remodeling of TGF-β/BMP signaling, shaping both inflammatory and differentiation-related responses. These findings point toward an integrated stress response, in which cytokine induction is coupled to suppression of canonical TGF-β signaling and activation of interferon-driven and pro-inflammatory programs, consistent with features of immunogenic cell death (34).

MIF expression increases during HNSCC progression, and serum levels are elevated in patients compared with healthy controls (35). Hypoxia induces cellular MIF expression and HIF-1α–dependent secretion (36), while ionizing radiation elevates MIF in cancer models (37). Consistently, we observed radiation-induced MIF secretion in Detroit-562 cells but not in FaDu or 2A3. Intracellular MIF was higher in HPV-negative tumors than in HPV-positive ones, yet *in vitro* conditioned medium from HPV-positive lines contained more MIF than from HPV-negative lines (7). Under hypoxia, HPV-negative lines released greater MIF than HPV-positive ones, likely reflecting higher baseline HIF-1α and constitutive secretion (7). In our study, Detroit-562 (HPV-negative) cells secreted substantially more MIF than 2A3, with hypoxia producing only modest increases. Differences with prior work may reflect hypoxia severity and duration (0.1% O_2_ for 48h vs. chronic 1% O_2_). Importantly, MIF also promotes neutrophil chemotaxis, underscoring its role in shaping the tumor microenvironment (38).

Elevated Serpin E1 levels correlate with poor prognosis in HNSCC, likely due to its role in promoting migration and inhibiting apoptosis (39). Both hypoxia and irradiation induce Serpin E1 expression in HNSCC, with hypoxia showing a stronger effect in FaDu cells (40). We similarly observed hypoxia as the dominant driver but also detected irradiation-induced Serpin E1 release in FaDu and 2A3, consistent with reports at higher radiation doses (41). *In vivo*, Serpin E1 correlated with local tumor control and was elevated in hypoxic xenografts (42). Clinically, baseline plasma Serpin E1 was significantly higher in HPV/p16-negative patients (10). Transcriptional levels of Serpin E1, together with MIF and VEGF, were increased in HNSCC lines under hypoxia. Interestingly, Serpin E1 expression returned to near baseline after reoxygenation, suggesting it as a candidate gene for the hypoxic tumor phenotype (43). Together, these findings indicate that Serpin E1 is modulated by both hypoxia and irradiation, and its elevated expression—particularly under hypoxic and HPV-negative conditions—may drive treatment resistance, supporting its potential as a prognostic biomarker and therapeutic target in stratified radiotherapy.

Our study is limited by the *in vitro* design, which may not fully capture the complexity of the *in vivo* tumor microenvironment. The use of a single hypoxic condition (1% O_2_) may not reflect the full spectrum of physiological hypoxia. While we analyzed key cytokines and apoptotic markers, other important non-apoptotic cell fates such as senescence or mitotic catastrophe were not evaluated. Finally, although we used representative HNSCC lines, the number per HPV group was limited, and broader validation in cell panels and *in vivo* models is needed to generalize these findings.

This study provides new mechanistic insights into how hypoxia modulates radiation response in HPV-positive versus HPV-negative HNSCC. Using a multi-layered approach—including proliferation, cell-cycle profiling, apoptosis markers, transcriptomics, and cytokine/chemokine secretion—we showed that hypoxia attenuated radiation-induced G2/M arrest and caspase-3 activation in a cell line–dependent manner and altered secretion of IL-8, MIF, and Serpin E1. These findings highlight reduced therapeutic efficacy under low-oxygen conditions and underscore the importance of integrating both HPV status and tumor hypoxia into biomarker-guided, stratified radiotherapy strategies for HNSCC. Together, these results provide a foundation for developing targeted interventions to overcome hypoxia-driven treatment resistance.

## Material and Methods

### Cell culture

The human cell line FaDu (305033, squamous cell carcinoma of the hypopharynx), and Detroit-562 (300399, metastatic cells of a pharyngeal carcinoma), were obtained from Cytion, Heidelberg, Germany. The 2A3 cell line (CRL-3212, ATCC, Manassas, VA, USA) was generated by transfecting FaDu cells with the E6 and E7 genes of HPV-16. Cells were cultured in RPMI medium (21875-091, Gibco, Grand Island, NY, USA) supplemented with 10% low-endotoxin fetal bovine serum (FBS, FB-1101/500, Biosera, France) and 1% penicillin/streptomycin (LM-A4118/100, Biosera, Nuaille, France). For 2A3 cells, penicillin/streptomycin was replaced with 0.2 mg/mL geneticin (10131-035, Gibco). Cell lines were authenticated via SRT by the vendor. Mycoplasma contamination tests were negative. Cells were cultured at 37°C in a humidified atmosphere under either normoxic (21% O_2_, 5% CO_2_) or hypoxic (1% O_2_, 5% CO_2_, 94% N_2_; InvivO_2_ 400, Baker Ruskinn, UK) conditions. All cell lines were cultured in parallel under normoxic and hypoxic conditions. Cells were maintained under their respective oxygen conditions continuously (FaDu and Detroit-562) or for seven days prior to experiments (2A3). Doubling time was calculated from non-irradiated controls cultured under ambient or hypoxic conditions.

### Conditioned medium

For conditioned medium collection, 2 × 10^5^ cells (FaDu, Detroit-562, and 2A3) were plated in duplicate in 6-well plates in 1.5 ml of RPMI medium supplemented with 10% FBS under either 21% or 1% O_2_. The following day, cells were irradiated with 0 or 6 Gy of gamma rays (dose rate: 1 Gy/min) using ^60^Co gamma-ray source (Chisostat, Chirana, Brno, Czech Republic). Hypoxic samples were placed in airtight pouches during irradiation. After irradiation, the medium was replaced with fresh oxygen-matched RPMI + 10% FBS, and cells were incubated under their respective oxygen conditions for 24 or 48 hours. At the designated time points, conditioned medium was aspirated (pooled from duplicates), centrifuged at 500×g for 10 minutes at 4°C to remove cell debris, and transferred to new tubes. Samples were processed under the corresponding oxygen conditions, aliquoted, and stored at −20°C until further use.

### Cell cycle assay

Cells were seeded and treated as described above. Twenty-four hours after irradiation, they were harvested separately under normoxic or hypoxic conditions using oxygen-matched buffers and trypsin. Cell cycle distribution was assessed using the Muse®Cell Cycle Kit (#MCH100106, Cytek, Fremont, CA, USA) according to the manufacturer’s protocol. Results are presented as the percentage of cells in G0/G1, S, or G2/M phases.

### Western blot

Cells were seeded and treated as shown above. Forty-eight hours after irradiation, they were harvested separately under normoxic or hypoxic conditions using oxygen-matched buffers and trypsin. The cells were processed as previously (44). Membranes were incubated overnight at 4 °C with primary antibodies: anti-β-actin (1:1 000; #3700, Cell Signaling Technology, Beverly, MA, USA), N-cadherin (1:1 000; #610921, BD, Franklin Lakes, NJ USA), vimentin (1:1 000; #GTX636980, GeneTex, Irvine, CA, USA), E-cadherin (1:500; #GTX636577 GeneTex) or anti-cleaved caspase-3 (1:1 000; #9664, Cell Signaling Technology). This was followed by incubation with corresponding HRP-linked secondary antibodies (anti-mouse IgG, #7076, 1:5 000; or anti-rabbit IgG, #7074, 1:2 000; both from Cell Signaling Technology) for 1 hour at room temperature. Protein bands were visualized using an enhanced chemiluminescent substrate (SuperSignal™ 34577/34095, Thermo Fisher Scientific). Optical densities of detected proteins were measured using ImageJ v1.52 (NIH, Bethesda, MD, USA).

### Cytokine array

Cytokine and chemokine profiling was performed using the Human Cytokine Array Kit (#ARY005B, Biotechne, R&D Systems, Minneapolis, MN, USA). We pooled conditioned medium from three independent experiments. Following manufacturer’s instructions, membranes were exposed in an Amersham A680 Imager (GE, Boston, MA, USA) for 1–20 minutes. Optical densities of detected cytokines and chemokines were quantified using ImageJ. Background-subtracted data were normalized to positive controls, log_10_-transformed, and z-scores were calculated.

### ELISA

Levels of IL-8, MIF, and Serpin E1 were quantified using an enzyme-linked immunosorbent assay (ELISA) kits (DY208-05, DY289 and DY1786, respectively; R&D system) according to the manufacturer’s instructions.

### RNA isolation, sequencing and analysis of differential expression

Total RNA was extracted from cells by TRI-Reagent (Sigma-Aldrich, St. Louis, MO, USA). Only RNA samples with an RNA integrity number (RIN) ≥7 determined by a Bioanalyzer (RNA 6000 Nano Kit, Agilent Technologies, Santa Clara, CA, USA) passed to library preparation. The TruSeq Stranded Total RNA LT Sample Prep Kit (Illumina, San Diego, CA, USA) was used to convert 0.5 μg of total RNA into a library of template molecules. The library was validated using Bioanalyzer (DNA 1000 Kit, Agilent) and quantified according to the manufacturer’s instructions by qPCR (KAPA Library Quantification Kit Illumina platforms, Kapa Biosystems, Wilmington, MA, USA) on a Quant studio 5 Real-Time PCR System (Thermo Fisher Scientific). Sequencing was performed on NextSeq 500 (Illumina). Low-quality reads were removed from the raw sequencing data, adaptor sequences clipped, and low-quality leading or trailing regions (below phred score 18) were trimmed using Trimmomatic v0.33 and BBDuk2 (both from the Joint Genome Institute, Walnut Creek, CA, USA). RNA sequencing data obtained from three biological replicates per group were aligned to the hg38 (45) reference genome using the HISAT (46) aligner. Reads mapped to gene regions were subsequently quantified with FeatureCounts (47), and differential expression analysis was performed using DESeq2(48). Genes were considered differentially expressed if they had an adjusted p-value (padj) <0.1. Genes which were evaluated as differentially expressed were visualized via InterctiVenn (49) and iDEP (50). WEB-based GEne SeT AnaLysis Toolkit 2024 (WebGestalt 2024), using Over-Representation Analysis based on Fisher’s exact test, was employed to analyze biological processes associated with differentially expressed genes (DEGs) (51). DEGs were identified from three independent RNA samples per line using thresholds of adjusted p<0.1 and absolute log_2_FC≥2.

### Statistics

Data are presented as arithmetic means ± standard deviations (SD). Statistical analysis was performed using two-way ANOVA followed by Tukey’s post hoc test. For comparisons between two independent groups, Student’s t-test was used. A p-value of <0.05 was considered statistically significant. Statistical analyses were conducted using GraphPad Prism v10.4 (GraphPad Software, USA, www.graphpad.com). Statistical methods and the number of independent experiments are provided in the respective figure legends.

## Acknowledgements

This study was funded by the Ministry of Education, Youth and Sports of the Czech Republic (MŠMT); project No. LUC23033. This publication is based upon work from the COST Action Converting molecular profiles of myeloid cells into biomarkers for inflammation and cancer (Mye-InfoBank), CA20117, supported by COST (European Cooperation in Science and Technology). Additional support was provided by Ministry of Health, Czech Republic - conceptual development of research organization (MMCI, 00209805) and by the project SALVAGE (P JAC; reg. no. CZ.02.01.01/00/22_008/0004644), co-funded by the European Union and the State Budget of the Czech Republic.

## Author Contributions

JP – conceptualization, designed and performed experiments, and wrote and revised the manuscript; FZK – designed and performed experiments, analyzed transcriptomic data, wrote and revised the manuscript; SV – designed and performed experiments, and revised the manuscript; OV – designed and performed experiments, and revised the manuscript; JH – designed and performed experiments, and revised the manuscript; RH – designed the study, performed experiments, secured funding and wrote and revised the manuscript; TP – designed the study, performed experiments, secured funding and wrote and revised the manuscript. All of the authors approved the final manuscript.

## Data Availability Statement

The NGS datasets generated and analyzed in this study will be made available in the NCBI public database.

## Supplementary Information

File: SI-Figures (PDF format)

Contains Figures S1–S4 showing transcriptomic and pathway analyses of HNSCC cell lines under hypoxia and gamma-irradiation, including volcano plots, KEGG pathway maps (Chemokines signaling and TGFβ), and ELISA data on cytokine/chemokine release 24 hours after irradiation.

## References

1. Hammond EM, Asselin MC, Forster D, O’Connor JP, Senra JM, Williams KJ. The meaning, measurement and modification of hypoxia in the laboratory and the clinic. Clin Oncol (R Coll Radiol). 2014;26(5):277–88.

2. Brown JM. Tumor hypoxia in cancer therapy. Methods Enzymol. 2007;435:297–321.

3. Hauth F, Toulany M, Zips D, Menegakis A. Cell-line dependent effects of hypoxia prior to irradiation in squamous cell carcinoma lines. Clin Transl Radiat Oncol. 2017;5:12–9.

4. Machiels JP, Rene Leemans C, Golusinski W, Grau C, Licitra L, Gregoire V, et al. Squamous cell carcinoma of the oral cavity, larynx, oropharynx and hypopharynx: EHNS-ESMO-ESTRO Clinical Practice Guidelines for diagnosis, treatment and follow-up. Ann Oncol. 2020;31(11):1462–75.

5. Borras JM, Barton M, Grau C, Corral J, Verhoeven R, Lemmens V, et al. The impact of cancer incidence and stage on optimal utilization of radiotherapy: Methodology of a population based analysis by the ESTRO-HERO project. Radiother Oncol. 2015;116(1):45–50.

6. Johnson DE, Burtness B, Leemans CR, Lui VWY, Bauman JE, Grandis JR. Head and neck squamous cell carcinoma. Nat Rev Dis Primers. 2020;6(1):92.

7. Kindt N, Descamps G, Lechien JR, Remmelink M, Colet JM, Wattiez R, et al. Involvement of HPV Infection in the Release of Macrophage Migration Inhibitory Factor in Head and Neck Squamous Cell Carcinoma. J Clin Med. 2019;8(1).

8. Rieckmann T, Tribius S, Grob TJ, Meyer F, Busch CJ, Petersen C, et al. HNSCC cell lines positive for HPV and p16 possess higher cellular radiosensitivity due to an impaired DSB repair capacity. Radiother Oncol. 2013;107(2):242–6.

9. Sorensen BS, Busk M, Olthof N, Speel EJ, Horsman MR, Alsner J, et al. Radiosensitivity and effect of hypoxia in HPV positive head and neck cancer cells. Radiother Oncol. 2013;108(3):500–5.

10. Brondum L, Eriksen JG, Singers Sorensen B, Mortensen LS, Toustrup K, Overgaard J, et al. Plasma proteins as prognostic biomarkers in radiotherapy treated head and neck cancer patients. Clin Transl Radiat Oncol. 2017;2:46–52.

11. Trellakis S, Bruderek K, Dumitru CA, Gholaman H, Gu X, Bankfalvi A, et al. Polymorphonuclear granulocytes in human head and neck cancer: enhanced inflammatory activity, modulation by cancer cells and expansion in advanced disease. Int J Cancer. 2011;129(9):2183–93.

12. Kalayci Yigin A, Azzawri A, Ozturk K, Cora T, Seven M. Determination of cytokine profile and associated genes of the signaling pathway in HNSCC. J Recept Signal Transduct Res. 2022;42(5):462–8.

13. Linkov F, Lisovich A, Yurkovetsky Z, Marrangoni A, Velikokhatnaya L, Nolen B, et al. Early detection of head and neck cancer: development of a novel screening tool using multiplexed immunobead-based biomarker profiling. Cancer Epidemiol Biomarkers Prev. 2007;16(1):102–7.

14. Shinohara Y, Washio J, Kobayashi Y, Abiko Y, Sasaki K, Takahashi N. Hypoxically cultured cells of oral squamous cell carcinoma increased their glucose metabolic activity under normoxic conditions. PLoS One. 2021;16(10):e0254966.

15. Cuisnier O, Serduc R, Lavieille JP, Longuet M, Reyt E, Riva C. Chronic hypoxia protects against gamma-irradiation-induced apoptosis by inducing bcl-2 up-regulation and inhibiting mitochondrial translocation and conformational change of bax protein. Int J Oncol. 2003;23(4):1033–41.

16. Yin X, Wei Z, Song C, Tang C, Xu W, Wang Y, et al. Metformin sensitizes hypoxia-induced gefitinib treatment resistance of HNSCC via cell cycle regulation and EMT reversal. Cancer Manag Res. 2018;10:5785–98.

17. Besso MJ, Bitto V, Koi L, Wijaya Hadiwikarta W, Conde-Lopez C, Euler-Lange R, et al. Transcriptomic and epigenetic landscape of nimorazole-enhanced radiochemotherapy in head and neck cancer. Radiother Oncol. 2024;199:110348.

18. Almatroodi SA, Alsahli MA, Almatroudi A, Verma AK, Aloliqi A, Allemailem KS, et al. Potential Therapeutic Targets of Quercetin, a Plant Flavonol, and Its Role in the Therapy of Various Types of Cancer through the Modulation of Various Cell Signaling Pathways. Molecules. 2021;26(5).

19. Kimple RJ, Smith MA, Blitzer GC, Torres AD, Martin JA, Yang RZ, et al. Enhanced radiation sensitivity in HPV-positive head and neck cancer. Cancer Res. 2013;73(15):4791–800.

20. Todorovic V, Prevc A, Zakelj MN, Savarin M, Brozic A, Groselj B, et al. Mechanisms of different response to ionizing irradiation in isogenic head and neck cancer cell lines. Radiat Oncol. 2019;14(1):214.

21. Arenz A, Ziemann F, Mayer C, Wittig A, Dreffke K, Preising S, et al. Increased radiosensitivity of HPV-positive head and neck cancer cell lines due to cell cycle dysregulation and induction of apoptosis. Strahlenther Onkol. 2014;190(9):839–46.

22. Gottgens EL, Bussink J, Leszczynska KB, Peters H, Span PN, Hammond EM. Inhibition of CDK4/CDK6 Enhances Radiosensitivity of HPV Negative Head and Neck Squamous Cell Carcinomas. Int J Radiat Oncol Biol Phys. 2019;105(3):548–58.

23. Jansen J, Vieten P, Pagliari F, Hanley R, Marafioti MG, Tirinato L, et al. A Novel Analysis Method for Evaluating the Interplay of Oxygen and Ionizing Radiation at the Gene Level. Front Genet. 2021;12:597635.

24. Ha J, Park M, Lee Y, Choi SH, Kim BS, Ha H, et al. AZD7648, a DNA-PKcs inhibitor, overcomes radioresistance in Hep3B xenografts and cells under tumor hypoxia. Am J Cancer Res. 2023;13(10):4918–30.

25. Moon SU, Choi SK, Kim HJ, Kumar Bokara K, Park KA, Lee WT, et al. The expression of human papillomavirus type 16 (HPV16 E7) induces cell cycle arrest and apoptosis in radiation and hypoxia resistant glioblastoma cells. Mol Med Rep. 2011;4(6):1247–53.

26. Ji Q, Li H, Cai Z, Yuan X, Pu X, Huang Y, et al. PYGL-mediated glucose metabolism reprogramming promotes EMT phenotype and metastasis of pancreatic cancer. Int J Biol Sci. 2023;19(6):1894–909.

27. Lee SH, Cho WJ, Najy AJ, Saliganan AD, Pham T, Rakowski J, et al. p62/SQSTM1-induced caspase-8 aggresomes are essential for ionizing radiation-mediated apoptosis. Cell Death Dis. 2021;12(11):997.

28. Sorensen BS, Busk M, Horsman MR, Alsner J, Overgaard J, Kyle AH, et al. Effect of radiation on cell proliferation and tumor hypoxia in HPV-positive head and neck cancer in vivo models. Anticancer Res. 2014;34(11):6297–304.

29. Gottgens EL, Bussink J, Ansems M, Hammond EM, Span PN. AKT inhibition as a strategy for targeting hypoxic HPV-positive HNSCC. Radiother Oncol. 2020;149:1–7.

30. Le QT, Fisher R, Oliner KS, Young RJ, Cao H, Kong C, et al. Prognostic and predictive significance of plasma HGF and IL-8 in a phase III trial of chemoradiation with or without tirapazamine in locoregionally advanced head and neck cancer. Clin Cancer Res. 2012;18(6):1798–807.

31. Gehrke T, Hackenberg S, Polat B, Wohlleben G, Hagen R, Kleinsasser N, et al. Combination of salinomycin and radiation effectively eliminates head and neck squamous cell carcinoma cells in vitro. Oncol Rep. 2018;39(4):1991–8.

32. Meidenbauer J, Wachter M, Schulz SR, Mostafa N, Zulch L, Frey B, et al. Inhibition of ATM or ATR in combination with hypo-fractionated radiotherapy leads to a different immunophenotype on transcript and protein level in HNSCC. Front Oncol. 2024;14:1460150.

33. Byers LA, Holsinger FC, Kies MS, William WN, El-Naggar AK, Lee JJ, et al. Serum signature of hypoxia-regulated factors is associated with progression after induction therapy in head and neck squamous cell cancer. Mol Cancer Ther. 2010;9(6):1755–63.

34. Kondoh N, Mizuno-Kamiya M. The Role of Immune Modulatory Cytokines in the Tumor Microenvironments of Head and Neck Squamous Cell Carcinomas. Cancers (Basel). 2022;14(12).

35. Kindt N, Preillon J, Kaltner H, Gabius HJ, Chevalier D, Rodriguez A, et al. Macrophage migration inhibitory factor in head and neck squamous cell carcinoma: clinical and experimental studies. J Cancer Res Clin Oncol. 2013;139(5):727–37.

36. Liu L, Wang J, Wang Y, Chen L, Peng L, Bin Y, et al. Blocking the MIF-CD74 axis augments radiotherapy efficacy for brain metastasis in NSCLC via synergistically promoting microglia M1 polarization. J Exp Clin Cancer Res. 2024;43(1):128.

37. Gupta Y, Pasupuleti V, Du W, Welford SM. Macrophage Migration Inhibitory Factor Secretion Is Induced by Ionizing Radiation and Oxidative Stress in Cancer Cells. PLoS One. 2016;11(1):e0146482.

38. Dumitru CA, Gholaman H, Trellakis S, Bruderek K, Dominas N, Gu X, et al. Tumor-derived macrophage migration inhibitory factor modulates the biology of head and neck cancer cells via neutrophil activation. Int J Cancer. 2011;129(4):859–69.

39. Pavon MA, Arroyo-Solera I, Tellez-Gabriel M, Leon X, Viros D, Lopez M, et al. Enhanced cell migration and apoptosis resistance may underlie the association between high SERPINE1 expression and poor outcome in head and neck carcinoma patients. Oncotarget. 2015;6(30):29016–33.

40. Schilling D, Bayer C, Geurts-Moespot A, Sweep FC, Pruschy M, Mengele K, et al. Induction of plasminogen activator inhibitor type-1 (PAI-1) by hypoxia and irradiation in human head and neck carcinoma cell lines. BMC Cancer. 2007;7:143.

41. Artman T, Schilling D, Gnann J, Molls M, Multhoff G, Bayer C. Irradiation-induced regulation of plasminogen activator inhibitor type-1 and vascular endothelial growth factor in six human squamous cell carcinoma lines of the head and neck. Int J Radiat Oncol Biol Phys. 2010;76(2):574–82.

42. Bayer C, Schilling D, Hoetzel J, Egermann HP, Zips D, Yaromina A, et al. PAI-1 levels predict response to fractionated irradiation in 10 human squamous cell carcinoma lines of the head and neck. Radiother Oncol. 2008;86(3):361–8.

43. Koong AC, Denko NC, Hudson KM, Schindler C, Swiersz L, Koch C, et al. Candidate genes for the hypoxic tumor phenotype. Cancer Res. 2000;60(4):883–7.

44. Perecko T, Pereckova J, Hoferova Z, Falk M. Cell-type specific anti-cancerous effects of nitro-oleic acid and its combination with gamma irradiation. Biol Chem. 2024;405(3):177–87.

45. Yates AD, Achuthan P, Akanni W, Allen J, Allen J, Alvarez-Jarreta J, et al. Ensembl 2020. Nucleic Acids Res. 2020;48(D1):D682-D8.

46. Kim D, Paggi JM, Park C, Bennett C, Salzberg SL. Graph-based genome alignment and genotyping with HISAT2 and HISAT-genotype. Nat Biotechnol. 2019;37(8):907–15.

47. Liao Y, Smyth GK, Shi W. featureCounts: an efficient general purpose program for assigning sequence reads to genomic features. Bioinformatics. 2014;30(7):923–30.

48. Love MI, Huber W, Anders S. Moderated estimation of fold change and dispersion for RNA-seq data with DESeq2. Genome Biol. 2014;15(12):550.

49. Heberle H, Meirelles GV, da Silva FR, Telles GP, Minghim R. InteractiVenn: a web-based tool for the analysis of sets through Venn diagrams. BMC Bioinformatics. 2015;16(1):169.

50. Ge SX, Son EW, Yao R. iDEP: an integrated web application for differential expression and pathway analysis of RNA-Seq data. BMC Bioinformatics. 2018;19(1):534.

51. Elizarraras JM, Liao Y, Shi Z, Zhu Q, Pico AR, Zhang B. WebGestalt 2024: faster gene set analysis and new support for metabolomics and multi-omics. Nucleic Acids Res. 2024;52(W1):W415-W21.

