## Supplemental figures 1-4 for "HPV status and oxygen tension shape transcriptomic, inflammatory, and cell cycle responses in HNSCC treated with ionizing radiation"

Running title:
**Hypoxia and radiation response in head and neck cancer**

Jana Pereckova[1], Filip Zavadil Kokas[2], Simona Voznicova[3], Ondrej Vasicek[3], Jitka Holcakova[2], Roman Hrstka[2]*, Tomas Perecko[1]*

[1] Department of Cell Biology and Radiobiology, Institute of Biophysics of the Czech Academy of Sciences, Kralovopolska 135, 612 00 Brno, Czech Republic
[2] Research Centre for Applied Molecular Oncology, Masaryk Memorial Cancer Institute, Zluty kopec 7, Brno, 656 53, Czech Republic
[3] Department of Biophysics of Immune System, Institute of Biophysics of the Czech Academy of Sciences, Kralovopolska 135, 612 00 Brno, Czech Republic

**Supplementary Information Files:**
Figure S1
Figure S2
Figure S3
Figure S4

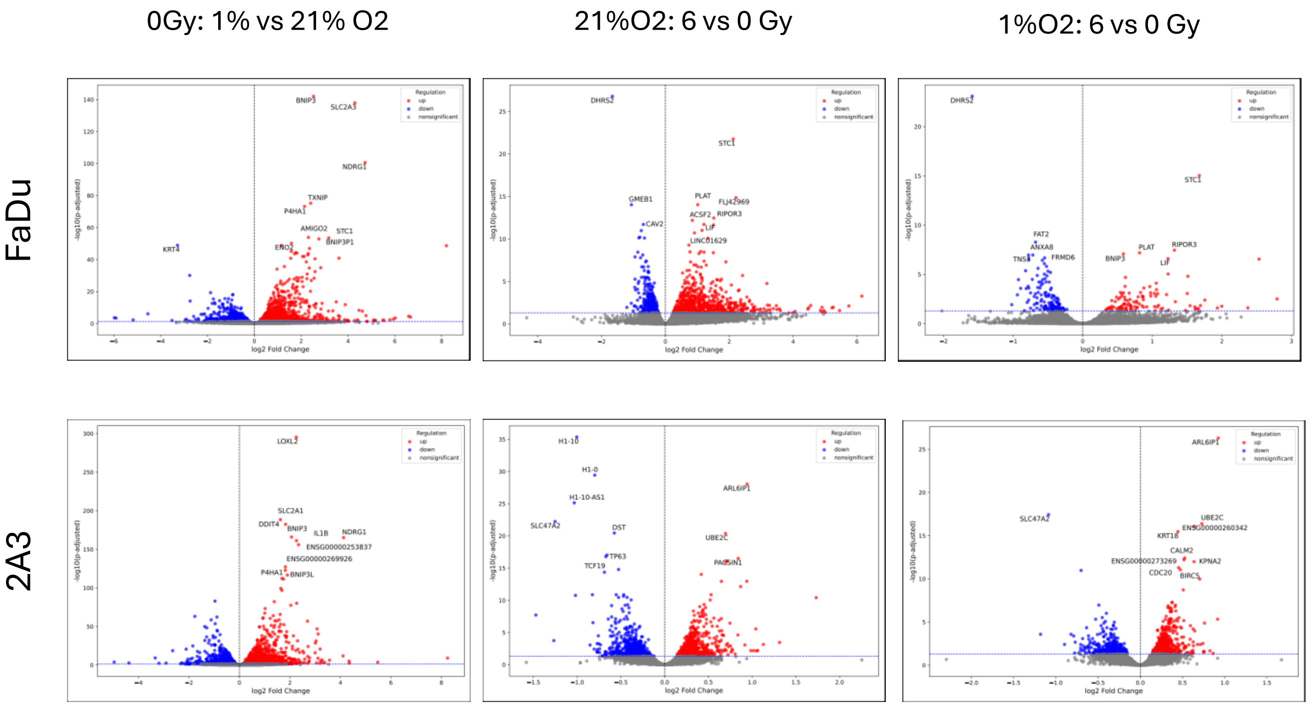


Figure S1. Effects of hypoxia and gamma-irradiation on differentially expressed genes in FaDu and 2A3 cell lines. Data are presented as log2 fold change versus negative decimal logarithm of the adjusted p-value. The ten most significantly altered genes are annotated. Threshold applied: adjusted p-value < 0.1.

FADU_0G1o_0G21o_Chemokine signaling

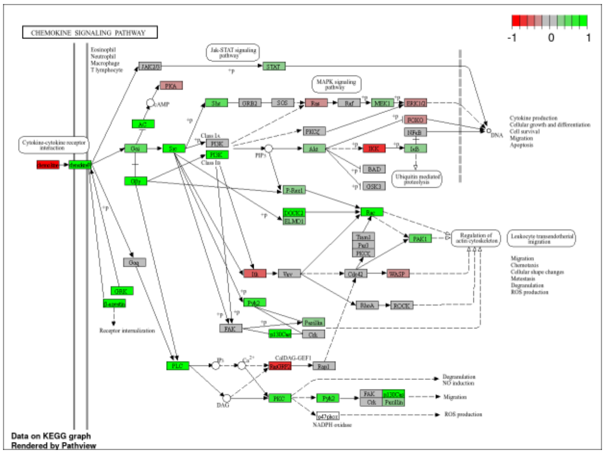


2A3_0G1o_0G21o_Chemokine signaling

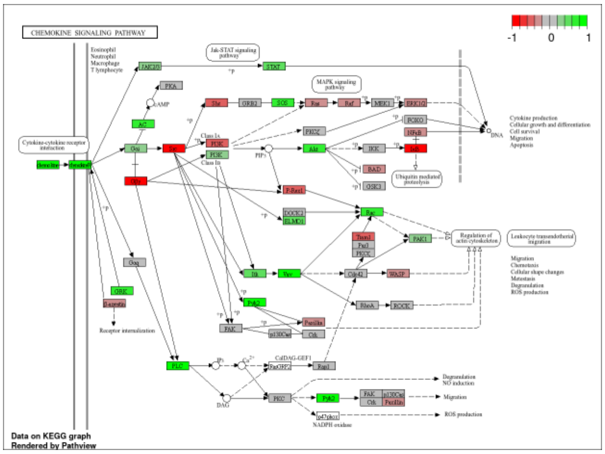


FADU_6G21o_0G21o_Chemokine signaling

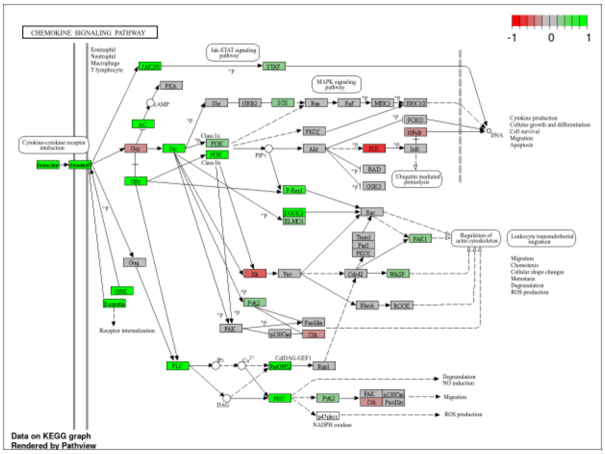


2A3_6G21o_0G21o_Chemokine signaling

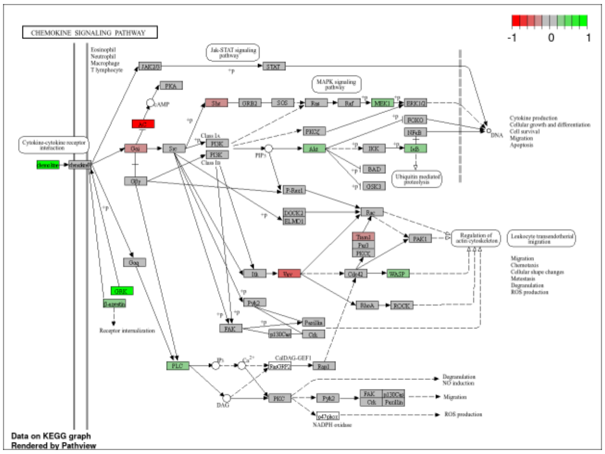


FADU_6G1o_0G1o_Chemokine signaling

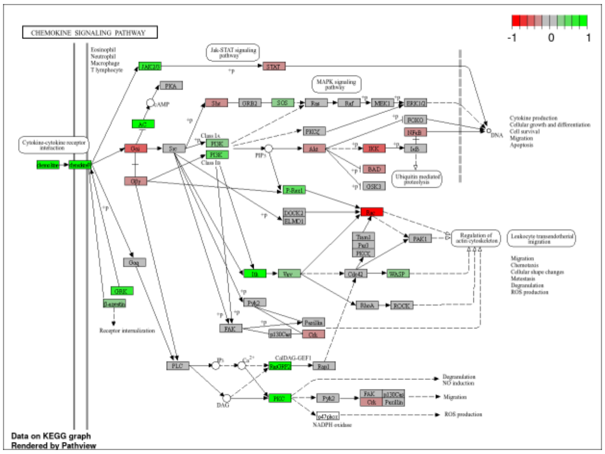


2A3_6G1o_0G1o_Chemokine signaling

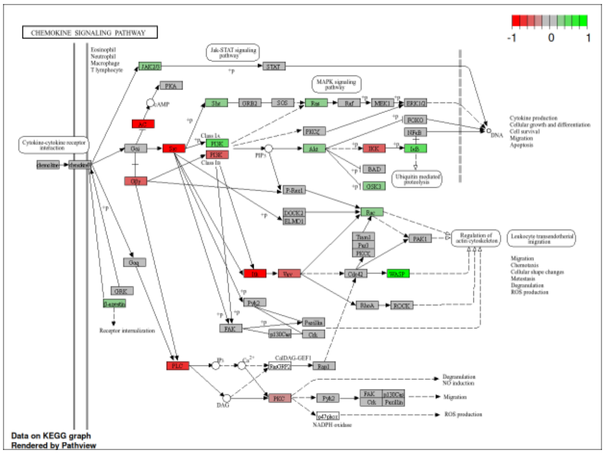


Figure S2. KEGG chemokine signaling pathway with mapped differentially expressed genes from FaDu and 2A3 cell lines after hypoxia and gamma-irradiation. Genes are color-coded according to log2 fold change: red indicates downregulation, green indicates upregulation, and grey denotes no significant change. Threshold applied: adjusted p-value < 0.1. The map was generated using the KEGG database and visualized with Pathview.

FADU_0G1o_0G21o_TGF

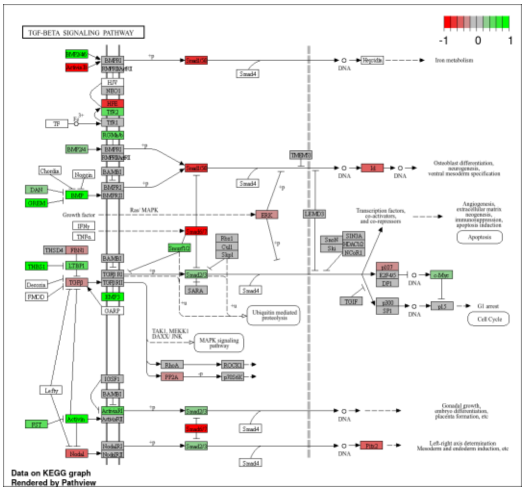


2A3_0G1o_0G21o_TGF

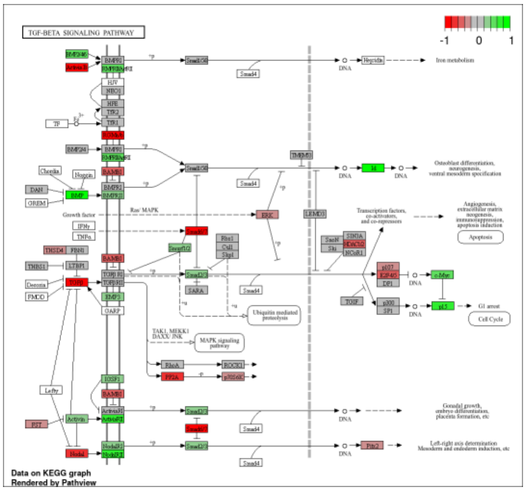


FADU_6G21o_0G21o_TGF

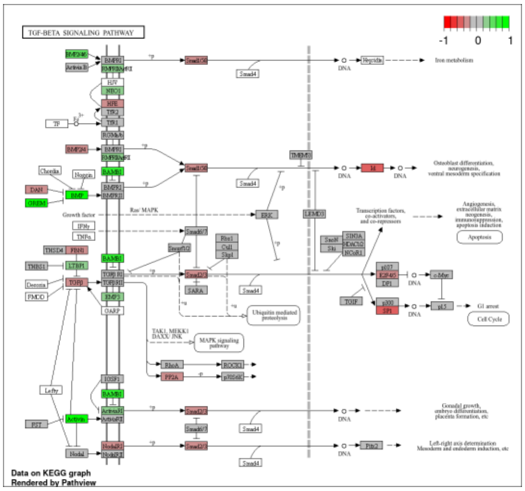


2A3_6G21o_0G21o_TGF

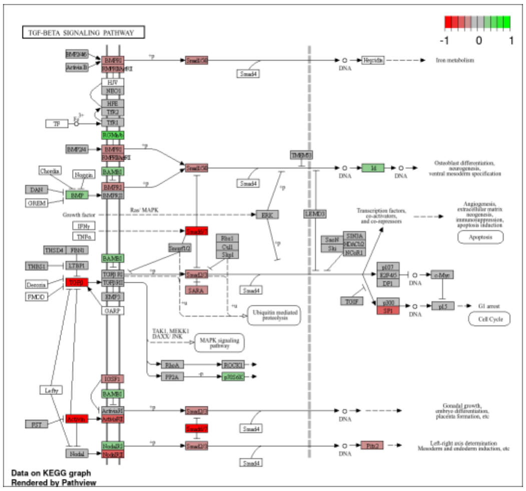


FADU_6G1o_0G1o_TGF

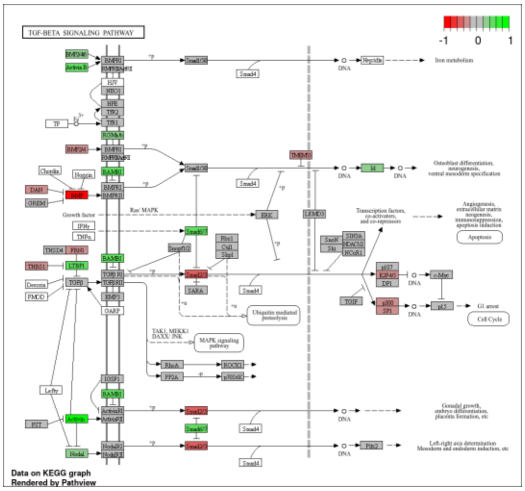


2A3_6G1o_0G1o_TGF

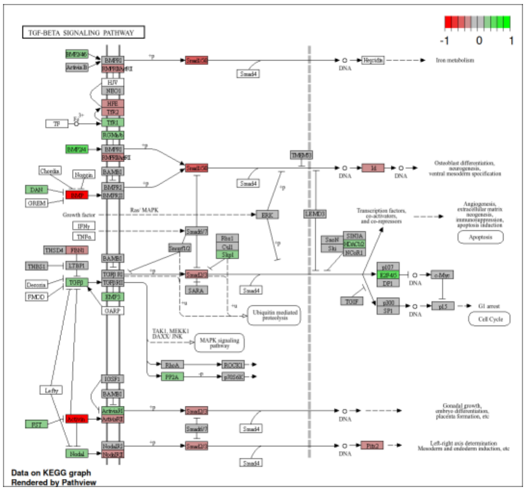


Figure S3. KEGG TGF-beta signaling pathway with mapped differentially expressed genes from FaDu and 2A3 cell lines after hypoxia and gamma-irradiation. Genes are color-coded according to log2 fold change: red indicates downregulation, green indicates upregulation, and grey denotes no significant change. Threshold applied: adjusted p-value < 0.1. The map was generated using the KEGG database and visualized with Pathview.

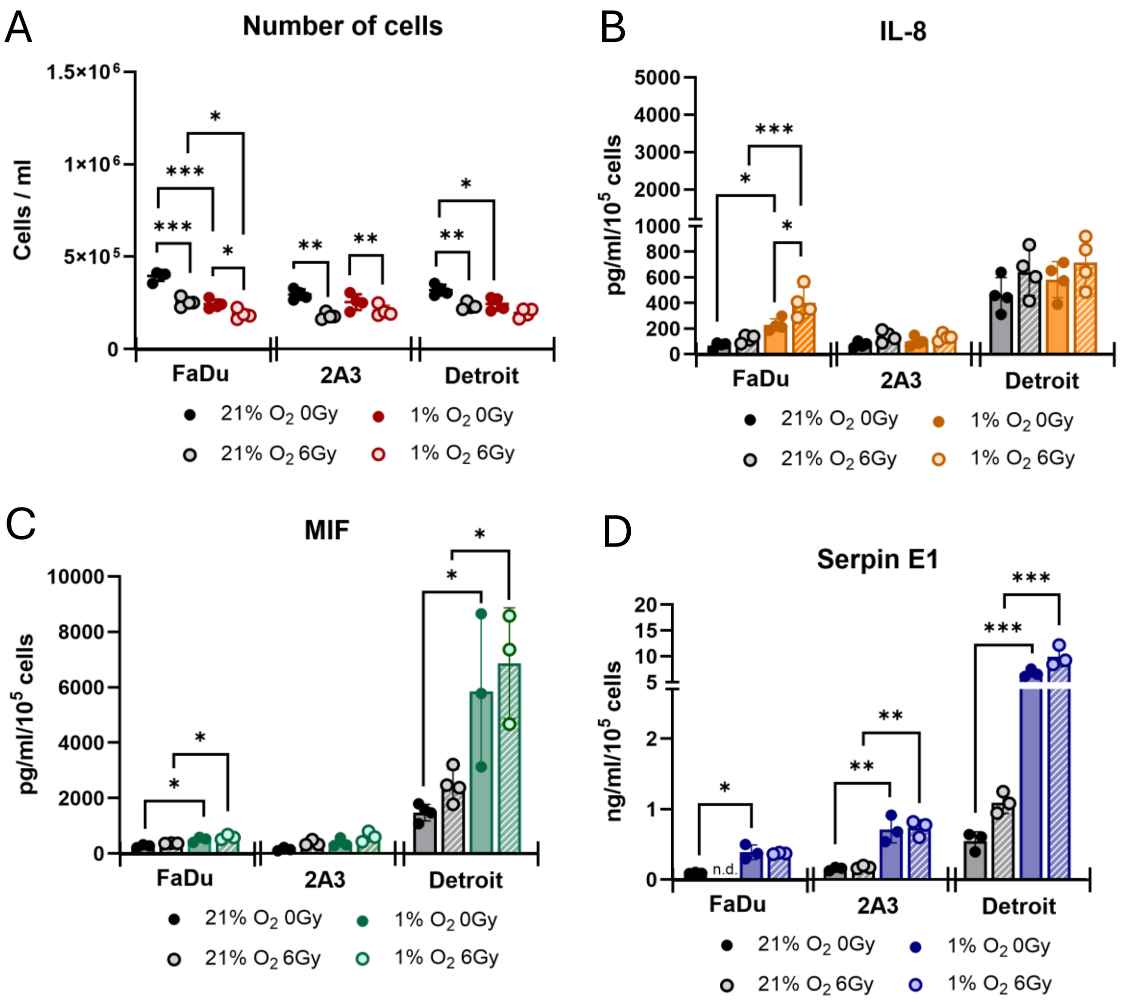


Figure S4. Effects of hypoxia and gamma-irradiation on cytokine/chemokine release in HNSCC 24 hours after gamma irradiation. (A) Cell numbers of FaDu, 2A3, and Detroit-562 cultured under normoxic (21% $O_2$) or hypoxic (1% $O_2$) conditions, assessed 24 hours after 6 Gy gamma-irradiation. Data represent mean ± SD from four independent experiments (n = 4). Statistical analysis: two-way ANOVA with Tukey's multiple comparisons test; *$p < 0.05$, **$p < 0.01$, ***$p < 0.001$. Levels of IL-8 (B), MIF (C), and Serpin E1 (D) in supernatants from FaDu, 2A3, and Detroit-562 cells cultured under normoxic or hypoxic conditions, assessed 24 hours after gamma-irradiation. Concentrations were normalized to cell numbers per condition. Data represent mean ± SD from three to four independent experiments (n = 3–4). Statistical analysis: two-way ANOVA with Tukey's multiple comparisons test; *$p < 0.05$, **$p < 0.01$, ***$p < 0.001$.
